# Cell Painting-based bioactivity prediction boosts high-throughput screening hit-rates and compound diversity

**DOI:** 10.1101/2023.04.03.535328

**Authors:** Johan Fredin Haslum, Charles Lardeau, Johan Karlsson, Riku Turkki, Karl-Johan Leuchowius, Kevin Smith, Erik Müllers

## Abstract

Efficiently identifying bioactive compounds towards a target of interest remains a time- and resource-intensive task in early drug discovery. The ability to accurately predict bioactivity using morphological profiles has the potential to rationalize the process, enabling smaller screens of focused compound sets.

Towards this goal, we explored the application of deep learning with Cell Painting, a high-content image-based assay, for compound bioactivity prediction in early drug screening. Combining Cell Painting data and unrefined single-concentration activity readouts from high-throughput screening (HTS) assays, we investigated to what degree morphological profiles could predict compound activity across a set of 140 unique assays.

We evaluated the performance of our models across different target classes, assay technologies, and disease areas. The predictive performance of the models was high, with a tendency for better predictions on cell-based assays and kinase targets. The average ROC-AUC was 0.744 with 62% of assays reaching ≥0.7, 30% reaching ≥0.8 and 7% reaching ≥0.9 average ROC-AUC, outperforming commonly used structure-based predictions in terms of predictive performance and compound structure diversity. In many cases, bioactivity prediction from Cell Painting data could be matched using brightfield images rather than multichannel fluorescence images. Experimental validation of our predictions in follow-up assays confirmed enrichment of active compounds.

Our results suggest that models trained on Cell Painting data can predict compound activity in a range of high-throughput screening assays robustly, even with relatively noisy HTS assay data. With our approach, enriched screening sets with higher hit rates and higher hit diversity can be selected, which could reduce the size of HTS campaigns and enable primary screening with more complex assays.

## Introduction

Drug discovery campaigns commonly rely on high throughput screens (HTS) to identify drug candidates with biological activity towards targets of interest. Such screens can involve probing millions of compounds. Although the throughput of these types of experiments has increased significantly thanks to technological advancements in automation and robotics, it is still a time and resource intensive process. Because of this, initial hit finding is generally done with simple assays such as biochemical assays to enrich the compound set before more resource-intense assays can be used further down the cascade. The initial screening assays are often very simple representations of the target biology and thus run the risk of producing false positive and negative results. Whereas false positives can be identified and removed by further probing with follow-up assays, false negatives can be problematic as they can filter out potentially interesting compounds. Thus, there is an interest in using as biologically relevant assays as possible early in the screening cascade.

One strategy to accelerate early hit finding is the use of computational methods to prioritize or select compounds deemed more likely to be active. Predicting bioactivity has been shown to enrich compound sets in HTS assays^1^. The allure of these types of approaches is a more efficient identification of drug candidates, which would reduce screening sets to only focus on the most relevant compounds. This, in turn, would enable earlier use of assays with higher biological relevance e.g., iPSC-derived or primary cells, which are typically restricted to later stage drug discovery due to cost and/or scarcity of biological material.

Today, bioactivity prediction is primarily performed using structure activity relation (SAR) models, although recently, alternative representations have been explored for compound property prediction ^2 3 4 5 6 7^. One such example is the use of phenotypic profiles^4^. This approach has emerged as an attractive alternative to SAR as they have proven to be capable of enriching compound sets while at the same time alleviating some of the drawbacks of structure-based models, such as low structural diversity and limited scaffold-hopping potential.

While it has been established that information present in phenotypic screens can be used to predict bioactivity in unrelated targets^4, 8^ these previous approaches relied on high-quality dose-response data as activity readouts. Crucially, in a real use-case setting, this information is unlikely to be sufficiently abundant at the early hit identification stage where the machine learning approaches proposed in these works are intended to be used.

In this work we explored a more practical setting that instead used single-concentration data readouts, which are more readily available, in combination with Cell Painting images which comes at a one-time upfront cost. We used a large-scale general purpose Cell Painting screen to capture phenotypic profiles of a library of available compounds and trained a model using small, focused bioactivity assay readouts for specific targets. The learned model could then make predictions on the rest of the compound library to identify which compounds to prioritize for screening (Figure 1a). The cost of the Cell Painting screen can be offset by reusing it in a similar manner to identify compounds for new sets of targets. This alternative approach has the potential to drastically reduce the number of assays and experiments needed in drug screening cascades. As only smaller compound sets are needed to be screened to train the predictive model for a particular target using this approach, assays of higher complexity and biological relevance could potentially be used.

**Figure 1.**
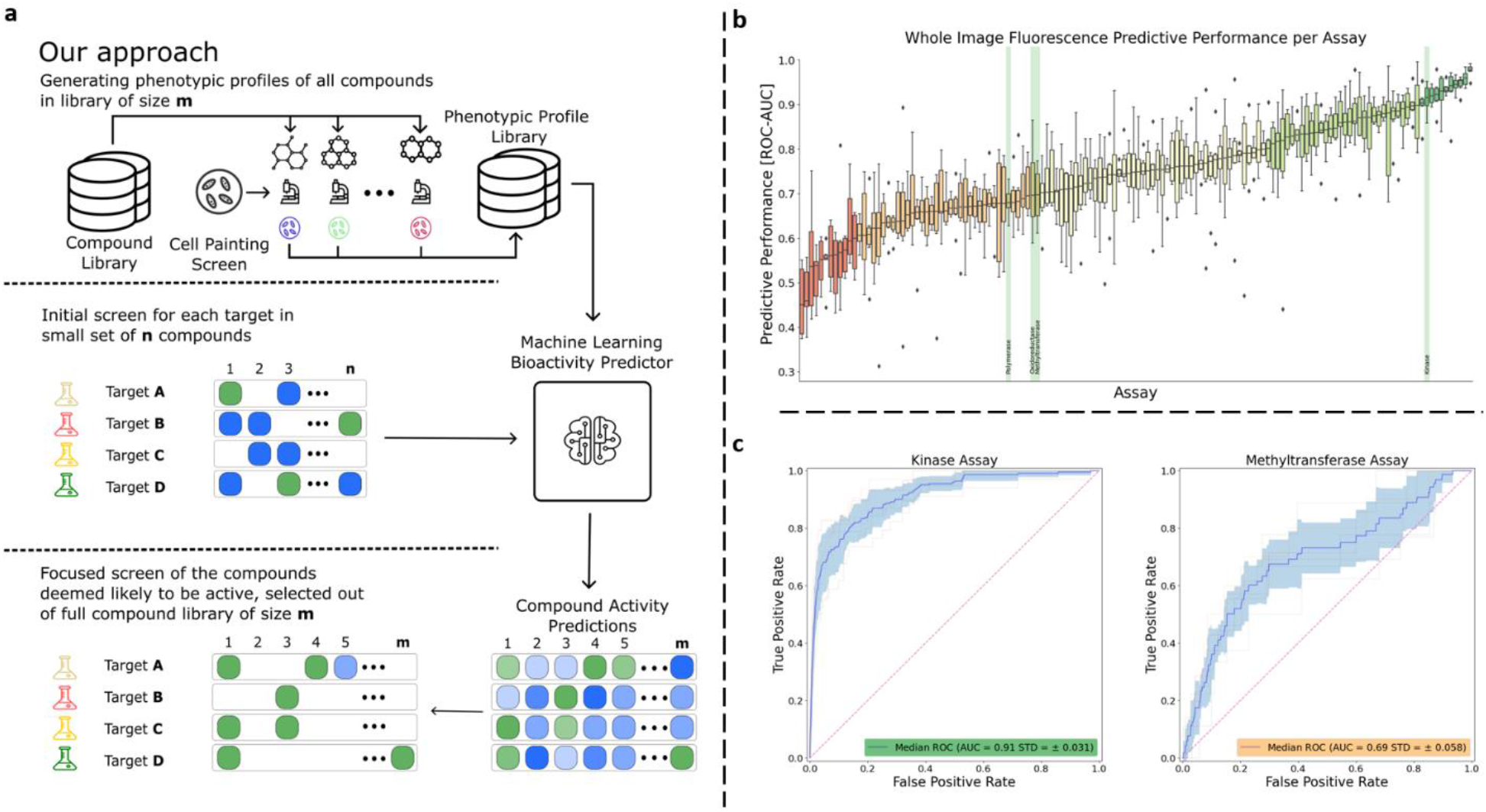
Using phenotypic screening for bioactivity prediction. A.) The envisioned approach utilizing phenotypic screening for bioactivity prediction. A single High Content Imaging screen using Cell Painting is used to generate phenotypic representations of each compound in the compound library m~10^6^. Small scale screens, n ~ 10^4^ for a target of interest are then used to generate activity readouts for a set of compounds used to train a machine learning model (green active, blue inactive). This model is then used to rank compounds for probability of activity and prioritize compounds for further screening (green predicted active, blue predicted inactive). B.) Box-plot of ROC-AUC performances of each assay in each cross-validation test-split, ranked in order of median ROC-AUC performance. Highlighted assays in green are investigated in secondary screens. C.) Receiver Operating Characteristic curve of two example assays. The dark blue line represents the average ROC-curve, the shaded area represents the standard-deviation intervals and the faded lines ROC-curves of individual cross-validation splits.

In addition, we also explored the use of different input modalities for bioactivity prediction. We considered fluorescence images, brightfield images, and image features extracted from the fluorescence images using classical image analysis approaches. We compared these image-based approaches to traditional structure-based approaches and demonstrated that compound bioactivity predicted from Cell Painting data outperformed the structure-based approaches across a wide range of different targets, target types, and assay technologies. These results were achievable even when working with noisy single point HTS readouts, as opposed to the more expensive dose-response data used in previous works. We found that image-based approaches outperformed structure-based models both in terms of predictive performance and structural diversity of the top ranked compounds.

Finally, we confirmed the validity of our predictions through a series of *in vitro* follow-up experiments which demonstrated that the bioactivity predictions of our models were consistent despite the inherent assay-to-assay variability. Taken together, our results demonstrated that phenotypic screening data can make bioactivity prediction more precise and efficient in a realistic drug screening scenario.

## Results

### Cell Painting-based phenotypic profiles are useful for bioactivity prediction

We selected a structurally diverse set of 8,300 compounds to be representative of a larger HTS screening library. This subset was screened in a Cell painting assay, an optimized high-content microscopy assay that utilizes a set of six fluorescent dyes to label different cellular components, including the nucleus, nucleoli, endoplasmic reticulum, mitochondria, cytoskeleton, Golgi apparatus, plasma membrane, actin filaments, as well as cytoplasmic and nucleolar RNA9. For each compound we also extracted corresponding single-point bioactivity data from the AstraZeneca HTS database. Together, the resulting dataset consisted of 8,300 compounds with Cell Painting images of compound-treated cells and associated single concentration bioactivity data in one or more of 140 unique assays, see Supplementary Figure 5 for more assay details. Each of those assays had at least 50 active compounds. The label matrix, according to compounds and assays, had a 47.8% fill rate and an average of 3% of the known compounds labelled as active (for more details of the data selection see Materials and Methods).

We split the data into six folds, with each compound appearing only in a single fold. Compounds that were structurally similar, based on ECFP-4 clustering, were assigned to the same fold to measure the ability of the model to identify actives in unknown regions of the compound space. This practice is commonly performed in Structure Activity Relationship (SAR) modeling. We applied cross-validation to the data, training a ResNet50 ^10^ in a supervised multi-task learning setup. The model was pretrained using ImageNet ^11^ and modified to accept 5-channel fluorescence images as input, and, given these inputs, trained to predict HTS bioactivity readouts for each of the 140 assays. In the cross-validation process, four of the data folds were used to train the model, one fold was used as validation to tune hyper-parameters, and one fold was used as a test set to evaluate performance (see Materials and Methods for training details).

The performance of the model, averaged across the six folds, was measured at a ROC-AUC of 0.744, but varied between the assays (Figure 1b). 62% of the assays achieved an ROC-AUC of 0.7 or more, which we deem ‘good performance’, while 30% reached 0.8 or higher (indicating ‘very good performance’), and a further 7% reached 0.9 or higher (indicating ‘excellent performance’). Overall, these results indicated that Cell Painting data possess valuable information related to bioactivity for a wide range of target and assay types, and that this relationship can be learned by a deep learning model using inexpensive single-concentration HTS activity readouts. We also investigated how the number of available activity readouts for a given assay influenced the ability of the model to make accurate predictions for that assay and found a weak positive correlation (Supplementary Figure 6).

### Comparing Cell Painting bioactivity predictions to other input modalities

As described above, we observed encouraging results using a multiplexed fluorescence Cell Painting screen to capture phenotypic profiles of a library of compounds. As it has been shown that brightfield images can be used to predict Cell painting features^12^, we wanted to investigate if the information content in the images would also be enough to predict bioactivity of compounds. Brightfield imaging has some advantages compared to Cell painting-stained cells as it can be performed on live cells and does not require staining of the cells and can be performed on simpler microscopes. These factors could significantly reduce the cost of the assay and enable kinetic assays, although potentially at the expense of less informative image data.

We also wanted to investigate whether the features learned by the neural network in our setup had an advantage over hand-crafted features. To this end, we extracted hand-crafted image features, hereafter referred to as Cell-Features, using the Columbus image-analysis software. Similar features could be extracted using free software such as Cell Profiler. We compared the performance of the neural network trained with Cell Painting images (‘whole image fluorescence’) versus a similar network trained with brightfield images (‘whole image brightfield’), as well as a neural network trained on hand-crafted ‘Cell-Features’. These image-based modalities were then compared against a standard structure-based approach using Extended Connectivity Fingerprints ^13^. Each model was assessed using the same cross-validation splits as described above. The fluorescence and brightfield images were used to train ResNet50 models, while the cell-features and structure-based data were used to train multi-layer perceptrons (See Materials and Methods for details).

Evaluation on the held-out test sets revealed that the predictive performance of the whole image fluorescence-based approach outperformed all other approaches (Figure 2.a). This approach reached an average ROC-AUC of 0.744 compared to the cell-feature based model at 0.726. Both fluorescence-based approaches (whole images and cell-features) outperformed the brightfield model at 0.704 ROC-AUC. The structure-based approach performed the worst at 0.686. Statistical analysis using Friedman rank sum test with the assay as blocking factor revealed significant performance differences (p = 4.47×10^-19^) between the modalities (Figure 2.a). Applying a post-hoc Nemenyi’s test, we find that the performance differences are significant between all modalities except for brightfield and structure. Although the brightfield image-based approach was outperformed by the fluorescence-based approach, it was still able to predict 49% of the assays with a ROC-AUC above 0.7 and even 5% above 0.9. This shows that the information captured in brightfield images can be linked to bioactivity in a wide range of assays and targets, which may justify using brightfield images in some cases despite their slightly inferior performance. We also tried to combine the brightfield images with the fluorescence images to see if it would offer any improvements; however, we saw no significant improvement in the predictive performance, indicating that the brightfield images contained little complementary information to the fluorescence images (data not shown).

**Figure 2.**
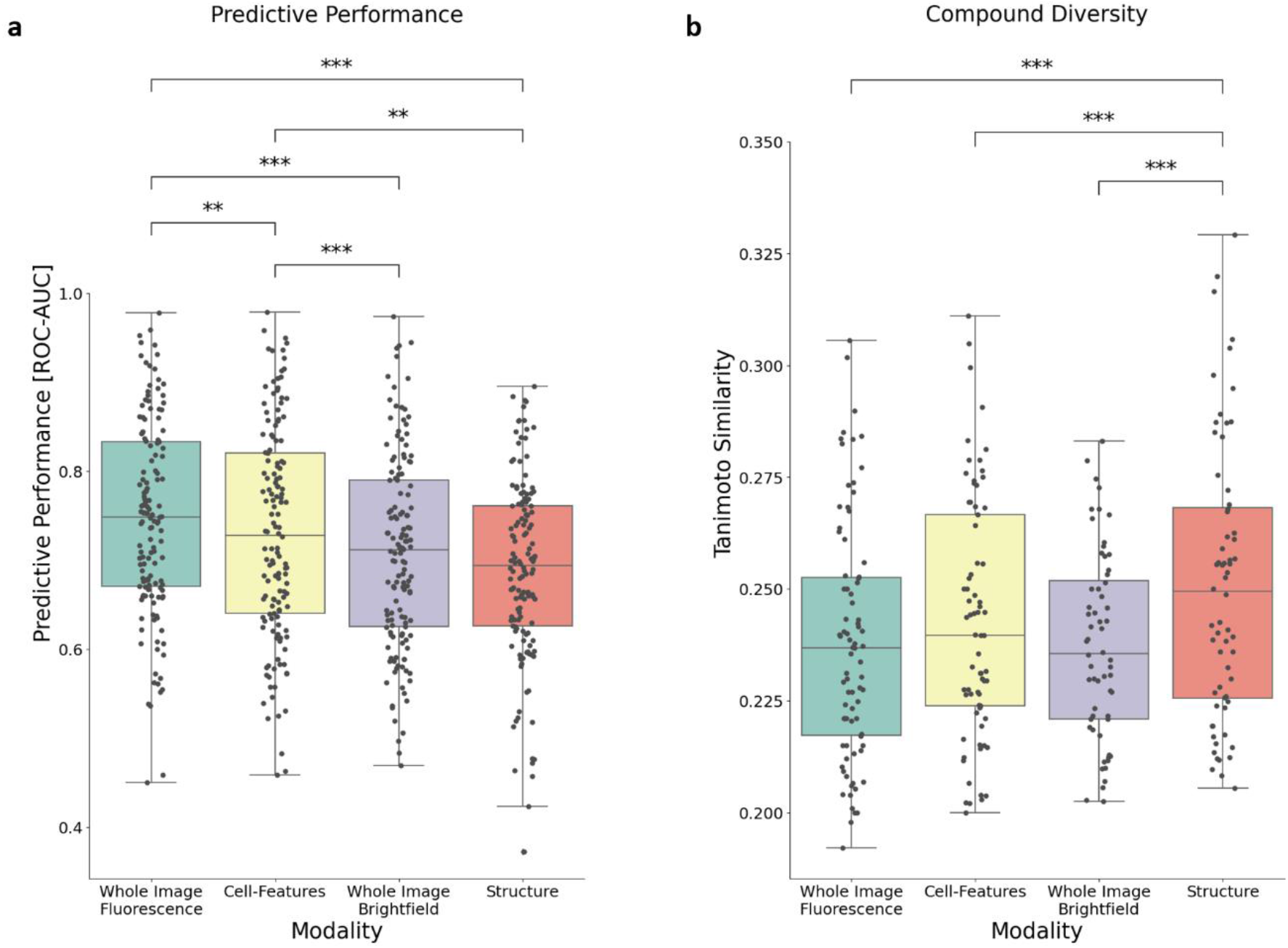
Variations in predictive accuracy and prediction diversity between input modalities. A.) Box plot of each modality type’s average ROC-AUC computed over each assay (n=140). Each dot represents the average ROC-AUC performance of an individual assay. B.) Box plot of average Tanimoto Similarity of top 20 ranked compounds to the closest known active in respective training set for each modality. Each dot represents the average Tanimoto similarity score per assay over all cross-validation splits. Only assays with performance above 0.6 ROC-AUC in all modalities were included (n=87). Statistical analysis was done using Nemenyi’s-Friedman post-hoc test, ** representing 10^-3^ < p < 10^-2^, *** representing 10^-4^ < p < 10^-3^.

A natural question to ask when predicting bioactivity of a compound library is: how chemically diverse are the top predictions? For each bioactivity prediction approach, we compared the structural diversity of the 20 top-ranked compounds to the known actives in the training set. Our analysis revealed that compounds predicted from images showed lower structural similarity, i.e. greater chemical diversity, than structure-based approaches. A Friedman rank sum test reveals a significant difference in the distribution of compound diversities (p=4.21×10^-12^). A Nemenyi’s post-hoc test showed that the chemical diversity of structure-based predictions was significantly lower than image-based predictions (Figure 2.b).

Overall, the Cell Painting fluorescence-based approach performed best both in terms of correctly predicting bioactivity (measured by ROC-AUC) and in terms of increased chemical diversity (measured by Tanimoto similarity). Because the Cell-Features approach also used the Cell Painting assay, it offered no cost savings yet performs worse than the Whole-Image Fluorescence approach. The Whole-Image Brightfield approach, however, may be an attractive alternative because the slight drop in predictive performance can be justified by other benefits compared to the Cell Painting assay.

### The impact of assay and target characteristics on performance

As seen in Figure 1b, the predictive performance of the Cell Painting fluorescence-image based model varied widely from assay to assay, ranging from 0.96 to 0.48 ROC-AUC. To gain a better understanding of which factors might impact the predictive capabilities of the model, we broke down the results to analyze performance attributed to various assay characteristics (Figure 3).

**Figure 3.**
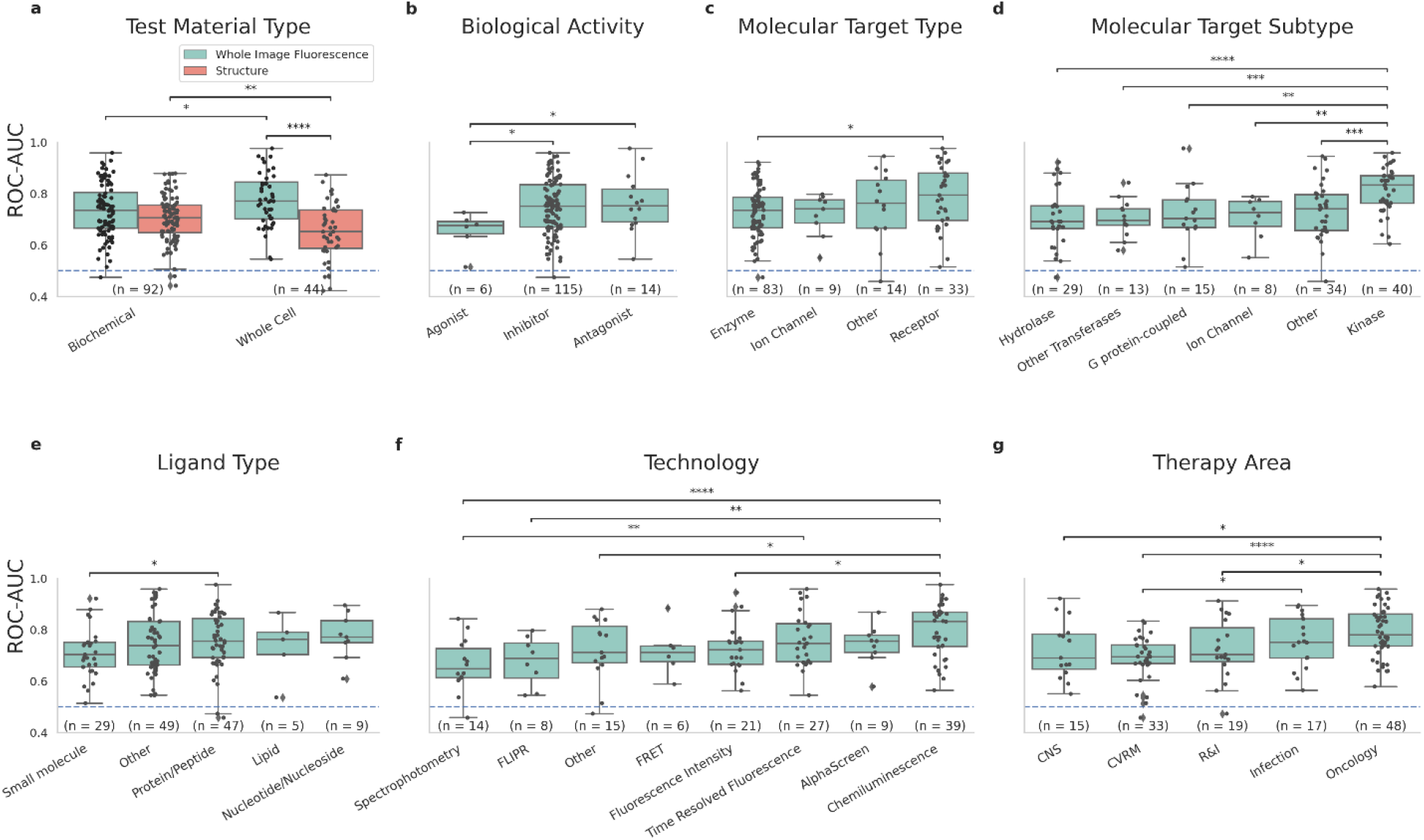
Predictive performance variations across assay characteristics. Predictive performance of various assays carried out in our experiments, grouped by assay characteristics. A.) The predictive performance of the image-based Fluorescence model compared to the structure based, when grouped by Test Material Type. B-G.) Comparison of Assay performance of the image-based Fluorescence model grouped by assay characteristics: B.) Biological Activity, C.) Molecular Target Type, D.) Molecular Target Subtype, E.) Ligand Type, F.) Assay Technology and G.) Therapy Area. R&I - Respiratory & Immunology, CVRM - Cardiovascular renal metabolism. Significance values calculated using non-parametric-ANOVA, * representing 10^-2^ < p < 5*10^-2^, ** representing 10^-3^ < p < 10^-2^, *** representing 10^-4^ < p < 10^-3^, **** representing p < 10^-4^.

The general trend we found was that the predictive performance was consistently good for different assay types. Likewise, predictive performance was consistently good across target types, therapy areas, and assay technologies. However, there were a few notable observations: the fluorescence-based methods were better at predicting cell-based assays than biochemical assays, and the structure-based method was better at predicting biochemical assays than cell-based assays (Figure 3.a.). Among molecular target subtypes, kinase targets appeared to benefit the most from our Cell Painting-based approach, performing significantly better than other molecular target subtypes (Figure 3d). We speculate that the high performance on kinase targets could be because kinase inhibitors are notoriously promiscuous and could have effects on multiple cellular pathways, giving rise to stronger phenotypic responses which are easier for the model to identify. Looking closer at performance across assay technologies (Figure 3.f.), we saw that predictions were especially accurate in Chemiluminescence assays, and less so with Spectrophotometry. Among the therapy areas covered in our experiments, performance in oncology assays was significantly better than other areas, possibly due to a higher fraction of kinase targets in that therapy area.

### Follow-up assays validate fluorescence-based predictions

Previous studies that aim to predict bioactivity from image-based assays have often limited their analysis to a single primary assay. But biological and technical noise in the process can lead to inaccurate and potentially overoptimistic results when the experiment is repeated. We put the predictions of our Cell Painting-based model to the test by running secondary assays for the same targets. We selected follow-up assays that were used in the screening cascades as secondary assays to triage HTS hits, spanning different assay technologies and different target classes: a methyltransferase, a polymerase, an oxidoreductase, and a serine kinase. The follow-up assays were chosen to represent a range of performances in the primary HTS assay, from a low ROC-AUC of 0.68 up to a very high performance of 0.91. The assays selected for follow-up are marked in Figure 1b. In each of the selected follow-up assays, a ranked list of predicted bioactivities was produced by our model. The majority of the top-ranked 5% of compounds were randomly sampled and included in the follow-up assay, along with selection of compounds selected uniformly at random (at least 500 compounds in total). The ROC-AUC reported in our experiments is computed using only the randomly selected compounds in order to keep the values comparable with the primary assay. The top-ranked compounds were included in order to make meaningful measurements of enrichment as it was expected that few of the randomly selected compounds would be active, due to the low assay hit rates (i.e. out of 500 randomly selected compounds, only 5-10 were expected to be active).

The results are shown in Figure 4. We found the ROC-AUC values in the follow-up assays to be consistent with the values from the primary assays. In fact, three out of the four follow-up assays performed slightly *better* than their respective primary assays (Figure 4a). Our results suggested that not only do the model predictions carry over to follow-up experiments, but that the expected range of performance is consistent as well.

**Figure 4.**
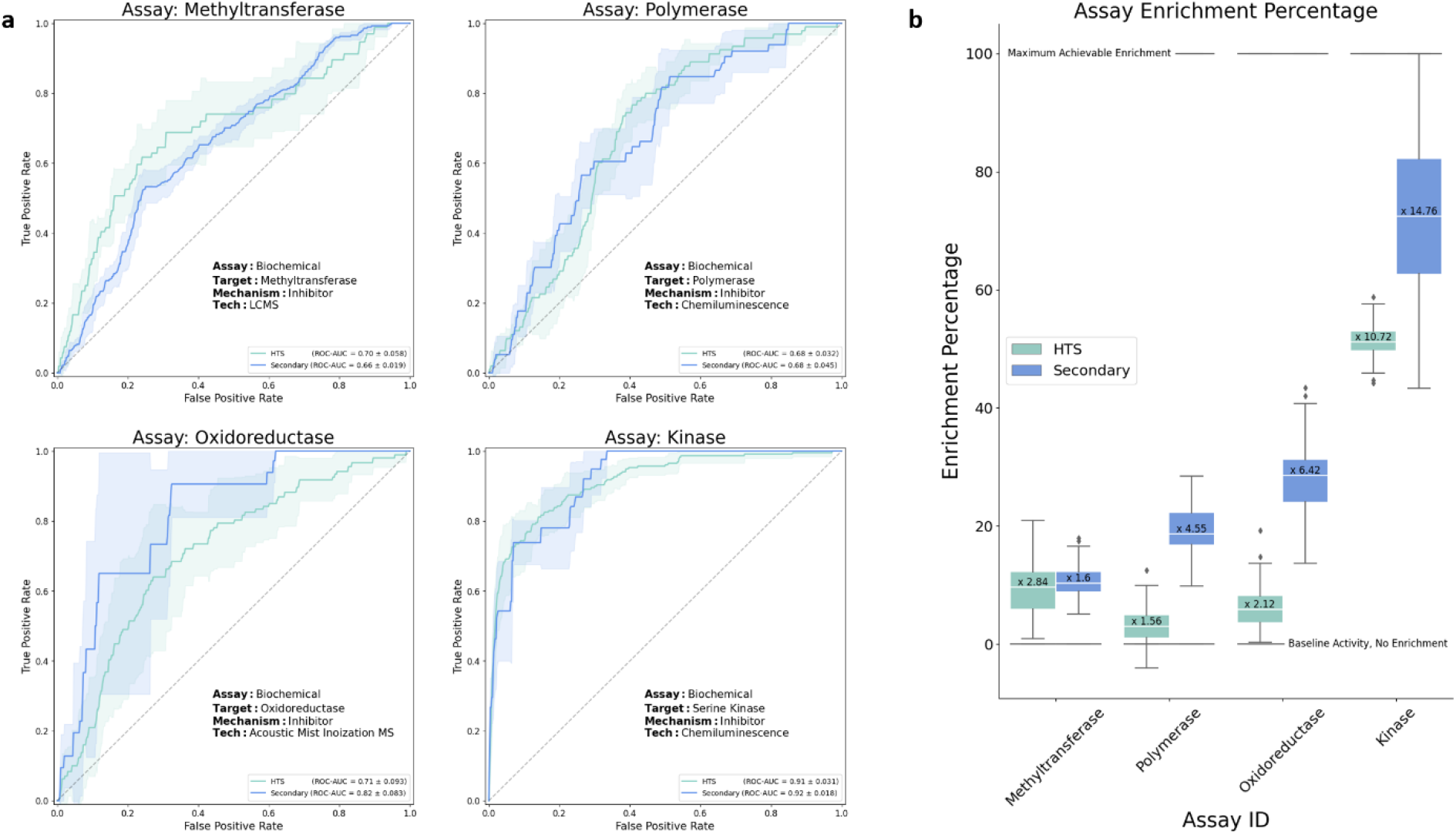
Validating model prediction in secondary screen. Model validation using follow-up screens in four of the assays. Top ranked compounds suggested from the fluorescence image-based models were screened in corresponding secondary assays (see text for full description). A.) Receiver operating curves for the four assays, primary HTS (green) and follow-up (blue), shaded area represent standard-deviation interval. ROC-AUC values for each of the four assays were calculated using the randomly sampled subset of compounds, green showing the average performance in the original HTS assays and blue representing the performance when using secondary screen activity readouts. B.) Enrichment values of the top 5% predicted compounds for each of the four assays. Enrichment was calculated for the HTS (green) and secondary (blue) activity readouts.

While ROC-AUC helps understand the predictive performance of the model, ultimately, what we care about is how the model can be used to enrich the compound sets. To calculate enrichment, using bootstrapping we probe the top 5% of ranked compounds for each assay (accordingly, the theoretical maximum enrichment from this experiment is 20x). The results appear in Figure 4b. Overall, the follow-up assays showed enrichment values in line with the primary assay, often better. The serine kinase assay, which performed well in the primary screen (ROC-AUC 0.91) showed an astoundingly high enrichment of 14x in the follow-up, which represents a significant improvement and suggests this assay could focus on a small, highly targeted set of compounds. The other assays, Oxidoreductase, Polymerase, and Methyltransferase, all had follow-up enrichments in line with the primary assay (6.4x, 4.6x, and 1.6x respectively). The follow-ups for Oxidoreductase and Polymerase showed higher enrichment than the primary assay, while the follow-up for Methyltransferase showed a slightly lower enrichment. However, it should be noted that the ROC-AUC was consistent between the primary and secondary. The weaker enrichment may be explained by the high baseline bioactivity in the Methyltransferase follow up assay, which limited the theoretical maximum enrichment of this assay to approximately 6x.

Several of the top ranked compounds were shown to be highly potent (low nM potency), indicating the quality of our predicted hits was high. Moreover, several compounds were predicted to be active by the model and were confirmed to be active, despite them having been labeled as inactive in the original HTS (data not shown). Their activity could be further confirmed in orthogonal assays, highlighting the robustness of our predictions and indicating that our model offers opportunities to rescue false-negative compounds. In summary, the assays probed in our follow-up experiments showed that the model performance was conserved through replication, even perhaps slightly better than what was expected. This indicated that the model’s predictions were driven by the targets and phenotypes, and not significantly affected by biases and noise in the assays. Furthermore, the enrichment levels we observed were high enough to reduce cost and speed up the screening process by filtering *in silico* compounds according to the ones the model predicts to be active.

## Discussion

Lead-identification in early drug-discovery is time and resource intensive, often relying on target-specific HTS campaigns to identify diverse, bioactive compounds. We assessed the possibility of rationalizing the lead-identification process using phenotypic-based bioactivity prediction models, trained with unrefined single-point concentration data, as opposed to more expensive dose-response data used in prior studies.

Our results show that a model trained using phenotypic data from a single general-purpose Cell Painting screen can predict bioactivity in a wide range of assays, outperforming commonly used SAR models in terms of both predictive performance and structure diversity. We validated the model’s performance in follow-up experiments in secondary assays. The results showed that Cell Painting-based bioactivity prediction using morphological profiles was feasible for a wide range of targets. Our approach has the potential to reduce the number of assays and experiments needed in drug screening cascades, which could allow for early screening in focused compound sets with assays of higher biological relevance. In addition, our results show that our brightfield-based model can perform slightly better than structure-based predictions, while also identifying more diverse compounds. This is in line with previous work showing that much of the information content of fluorescence images can be inferred from brightfield images ^12, 15, 16^ While the predictive performance of brightfield images does not reach the level of fluorescence images, brightfield offers a cheaper alternative for phenotypic profiling. Moreover, the use of brightfield images also opens to the possibility of live cell imaging to include temporal information of compound effects in cells.

Previous work established the information link between phenotypic screening data and assay activity ^4 8^. Our results extend this by showing the capacity to learn from more readily available, but relatively noisy, unrefined single-point activity readouts. These results could open the possibility of significantly reducing compound sets screened in HTS or actively applying phenotypic-based bioactivity predictors in focused screens where only few activity readouts can be gathered, which cannot be scaled to HTS format. While bioactivity prediction models using compound structures as input has the advantage of requiring no *in vitro* data, alternative input representation separated from the molecular structure have been used successfully previously. The motivation for generating input data *in vitro* is that the proxy descriptor of a compound could potentially avoid the problems SAR models have with scaffold hopping and increasing the diversity amongst predicted hits. Previous work using activity-^5, 7^, transcriptomics-^14^ and phenotypic fingerprints^4, 8^ are examples of such approaches.

In this study, our models considered individual input-modalities separately. In future works, input modalities may be combined within a single model. Previous work combining compound structure information with activity fingerprints showed improved performance in bioactivity prediction^5^. The same was shown to be true for cell state, where gene-expression and phenotypic data provided complementary information boosting performance prediction^17^. Thus, we expect that such an approach would also boost the performance on bioactivity predictors.

Beyond the inclusion of auxiliary input modalities, performance can probably be improved by employing more powerful deep learning model types ^18, 19^, for example, state-of-the-art deep learning networks such as Vision-Transformers, which have showed improved performance in medical imaging tasks ^20^. Other training strategies such as Self-Supervised Learning has also the potential to further improve performance; however, we leave such exploration for future work. While phenotypic-based bioactivity prediction requires *in-vitro* data, such datasets do not need to be expensive to generate; the Cell Painting assay protocol was designed to be rich in information while low-cost to perform ^9^. Moreover, much of the predictive performance can be achieved using brightfield images, thus reducing the need for staining reagents and advanced microscopes. In a drug discovery setting, the initial expenses for a single Cell Painting HTS would be easily recouped, as traditional HTS assays could be reduced and replaced by bioactivity predictors using the phenotypic data generated, combined with smaller, focused screens. Once established, a general high-content dataset could be exploited for other use cases, such as to predict compound mode of action or toxicity^21 22 23^. The value it brings from multiple applications may justify the resources required to establish it.

In summary, we have shown that phenotypic screening data combined with readily available single concentration data can be used for bioactivity prediction in a wide range of assays, with high performance across different target classes, assay technologies, and disease areas. Beyond the use of fluorescence data, we have also shown that brightfield data can reach a performance level comparable to or better than structure-based predictors, highlighting that staining the cells might not be necessary, but that it does provide slightly better predictions. Overall, the results paint a positive picture for phenotypic-based bioactivity models to complement structure-based predictors, where data can be generated in a cost-effective manner and with several use-cases.

## Methods

### 1. Generation of the Cell Painting and HTS dataset

A set of 8,300 compounds was selected and screened in the Cell painting assay. The compounds set was selected based on chemical diversity, known annotations, compound availability, and on general representation in historical HTS screens. The Cell Painting staining procedure was performed according to the protocol by Bray et al.^9^ with some adjustments to stain concentrations and methodology as described recently^22^. Imaging was performed as described by Trapotsi et al^22^ imaging the five different fluorescent channels, as well as brightfield images at 3 focal planes of 4 μm distance.

Corresponding HTS activity data were extracted from an internal HTS assay database. Historical HTS data were annotated as binary active/inactive using thresholds individually determined for each assay during assay development. Data were not available for all the selected compounds in all available historical HTS screens and we set a threshold of a minimum number of 50 active and 50 inactive datapoints required for a screen to be included. The included screens covered a diverse range of assay technologies, target types, therapeutical areas, etc. Following these criteria, the resulting dataset included around 70,000 images, covering 8,300 compounds with associated activity data in 140 unique assays. The label matrix had a 47.8% fill rate and an average of 3% of the known compounds labelled as active.

### 2. Bio-activity prediction setup and evaluation

#### 2.1 Data splits and cross-validation

Using a cross-validation setup, we split the data into 6 different folds with each compound only included in one-fold. Using RDKit^24^, all compound SMILES representations were converted to ECFP4 1024 Bit, with salts removed.

Using RDKit *Butina ClusterData* function, the ECFP4 representations was used to group the data into unique clusters. These clusters were divided into 6 unique folds of similar size such that all structurally similar compounds, belonged to the same fold.

#### 2.2. Prediction setup

Depending on the input modality, different Machine Learning models were used. Multi-Layer Perceptrons (MLPs) were used for the cell-feature-based model and the structure-based, described below in 3.3 and 3.4. For the Fluorescence and Brightfield images ResNet50s were used.

All models were trained to predict if a compound was active or inactive in each of the unique 140 assays as a multi-label binary prediction task. All networks were trained with 140 output neurons representing the 140 unique assays. A sigmoid activation function was used to normalize the range of values for each output neuron individually to the range of [0,1].

The models were trained with Binary Cross-Entropy combined with Focal Loss ^25^. Both propagate a loss signal from each of the output neurons of the network. Given the fact that not all compounds have been tested in all assays, the label matrix is incomplete. Thus, the activity of many of the compounds are unknown and no loss signal backpropagated from those neurons. Area Under the Receiver Operating Characteristic Curve (ROC-AUC) was used to evaluate the model’s ability to separate the actives from inactive. Since many activity labels are missing, the performance is only calculated for compounds with known activity values. We report both the mean ROC-AUC over all assays as well as the individual ones.

### 3. Approach

#### 3.1 Fluorescence image-based model

Fluorescent microscopy images were stored as 16-bit TIFFs of size 1992×1992. The images were pre-processed and normalized such that the top and bottom 1 percentile intensity values were clipped for each image to remove noise and outliers.

Before being sent to the network as input during training, the images were augmented, including spatial down sampling, random cropping, horizontal and vertical flipping, random 90-degree rotations and colour shifting.

The augmented images were used as input to a ResNet50 model pre-trained on ImageNet and adapted to allow 5 channel input images by adding two channels to the input convolutional filter. The linear layer of the pre-trained model was replaced with a re-initialized one with 140 output neurons to match the number of assays.

All models were trained on two NVIDIA-Tesla 32Gb GPUs, using Pytorch DDP. A hyper-parameter search was performed using nested cross-validation in one of the cross-validation splits. The identified hyper-parameters were then used to fine-tune the pre-trained ResNet50, using early stopping based on the validation ROC-AUC performance. (Learning Rate 0.2/256, Optimizer SGD, weight decay 10^-4^, learning rate dropping on plateau). A model was trained for each of the Cross-Validation splits, with the best performing model based on validation-set performance being used to predict the likelihood of activity for the respective test set split..

#### 3.2 Brightfield image-based model

Brightfield microscopy images were stored as 16-bit TIFFs of size 1992×1992 with three images captured at different focal planes at each site (+/- 4 μm around the central Z plane). The images were pre-processed and normalized such that the top and bottom 1 percentile values were clipped for each image to remove noise and intensity outliers.

The Brightfield image-based model was trained following the same procedure as for the fluorescent image-based model, although the default setting of three channels as input was used, stacking the three focal planes into one image. Hyper-parameter tuning and evaluation were kept the same as for the fluorescent images.

#### 3.3. Cell-feature model

Single-cell image features were extracted for each plate using the Columbus software package (v 2.9.1, PerkinElmer). Features such as nucleus size, cell radius, average nuclei intensity, etc, were extracted. In total 1,176 features were used for each cell. This was done for each cell and averaged per well. The averaged data were normalized by z-normalization per plate using DMSO controls using feature-wise median and median absolute deviation. Features with a variance below 1.0 were deemed uninformative and removed.

Following feature normalization and removal, the remaining features were used as model input. A Multi-layer perceptron (MLP) was used for prediction. Similar to the previous two model types, binary-cross entropy combined with focal loss was used to train the model.

Due to the more manageable size and computational requirements of the feature-based model, a full hyper-parameter tuning was done using nested cross-validation in each of the cross-validation splits, searching for optimal, model-depth, layer-width, weight-decay and learning rate (3-hidden layers, 1024 layer width, weight decay 0.0, learning rate 5.0).

#### 3.4. Structure fingerprints model

Each of the 8,300 compounds were represented using the commonly employed Extended Connectivity Fingerprints (ECFP) with a compound component diameter of 4. Using RDKit^24^, all compound SMILES^26^ representations were converted to ECFP4 with salts removed.

A Multi-layer perceptron (MLP) was used for the predictions. Binary-cross entropy combined with focal loss was used to train the model.

Similar to the Cell-feature model, a full hyper-parameter tuning was done using nested cross-validation in each of the cross-validation splits. Searching for optimal, model-depth, layer-width, weight-decay and learning rate (3-hidden layers, 512 layer width, weight decay 0.0, learning rate 2.0).

### 4. Statistical methods

Performance variations between model types were analysed using Friedman rank sum test, using the assays as blocking factors. This was calculated using Friedman-Chi-Square test using SciPy^27^ stats package followed by Nemenyi’s-Friedman post-hoc test.

Performance differences depending on assay characteristics was analysed using one-way ANOVA with post-hoc tests, calculated using Kruskal-Wallis test in SciPy stats package, followed by Conover pair-wise test to determine if there were any statistically significant differences between sub-groups.

### 5. Diversity evaluation

Tanimoto Similarity^28^ between the ECFP4 fingerprints of compounds was used to determine how structurally similar each compound pair was. To assess the diversity of the top ranked compounds according to each predictive model, the top 20 ranked compounds in each test set were compared to the known actives in their respective training set. Each top ranked compound was compared to all the known actives and the most similar one was identified for each of the 20 compounds, meaning the one with highest Tanimoto Similarity was then assigned as the most similar.

### 6. Follow-up screening and calculation of enrichment

Top ranked compounds in four of the assays were selected for follow-up validation in secondary screening. These compounds were selected based on their activity scores in the test set. This allowed us to select the top compounds from the full dataset without data leakage. The Fluorescence Whole Image based model type was used to assign activity scores for each compound. The top ranked five percent of compounds were randomly sampled for each of the four follow-up assays, with varying numbers of compounds selected for each of the assays. In addition, a random subset of compounds was also sampled for follow-up screening and used to calculate ROC-AUC metrics in the follow-up assays.

In each of the available follow-up assays a baseline estimate of assay activity was established by probing 600 randomly sampled compounds. This gives an estimate of the overall hit rate in each assay.

The likelihood of activity for each of the compounds in all six test-splits were combined and ranked together. The compounds deemed most likely to be active according to the Cell Painting Fluorescent Whole Image ResNet50 model were screened.

The ROC-AUC values in the follow-up screen were then evaluated using the randomly sampled compounds. The randomly sampled compounds were also used to assess the baseline hit rate for each of the assays, which was used for the enrichment analysis. The enrichments at different percentiles were then calculated using bootstrapping of the activity values of the compounds above that percentile.

## Acknowledgements

The authors would like to thank the AZ SCP team for assisting with computational resources. We also thank Eva Hansson, Craig Hughes, Linda Fredlund, Fredrik Wågberg, Nancy Su, Vijay Chandrasekar, Xiang Zhai for their assistance in generating the imaging dataset and or running wet-lab experiments.

## Contributions

J.F-H., J.K. C.L., KJ.L., R.T., E.M. and K.S. performed study concept and design, J.F-H. performed the computational implementation and analysis. KJ.L. and E.M. conducted wet-lab work. All the authors contributed to the writing of the paper and have read and approved the current version of the paper.

## Data availability

The HTS datasets generated and analysed in this study are not publicly available due to them being AstraZeneca proprietary information.

## Code availability

The code can be made available upon reasonable request to the authors.

## Supplementary figures

**Figure 5 – Supplementary figure.**
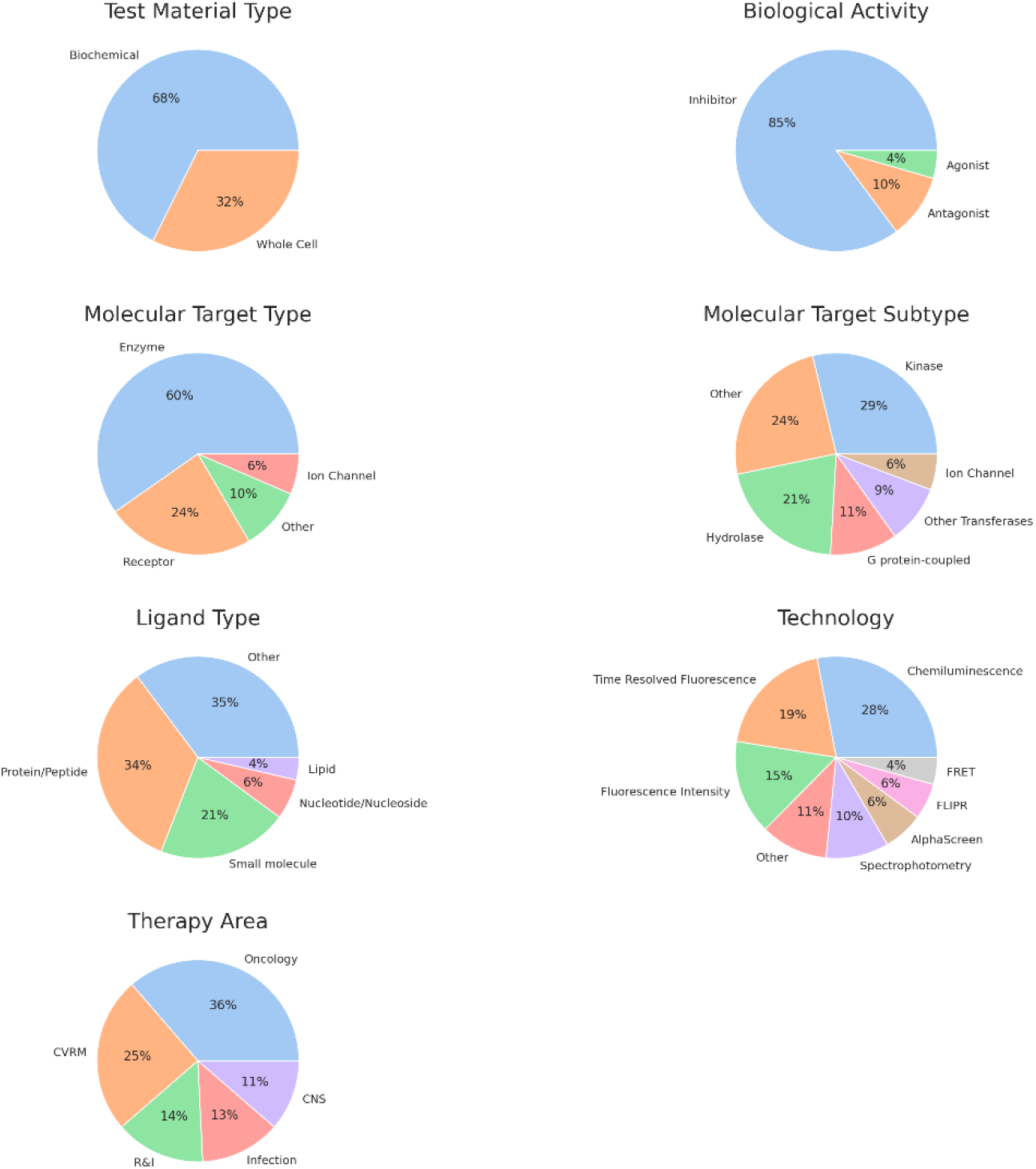
HTS assay characteristics. Pie-chars of assay characteristics of the HTS assay used. Grouped by major classes, rare characteristics added into “other” group.

**Figure 6 – Supplementary figure.**
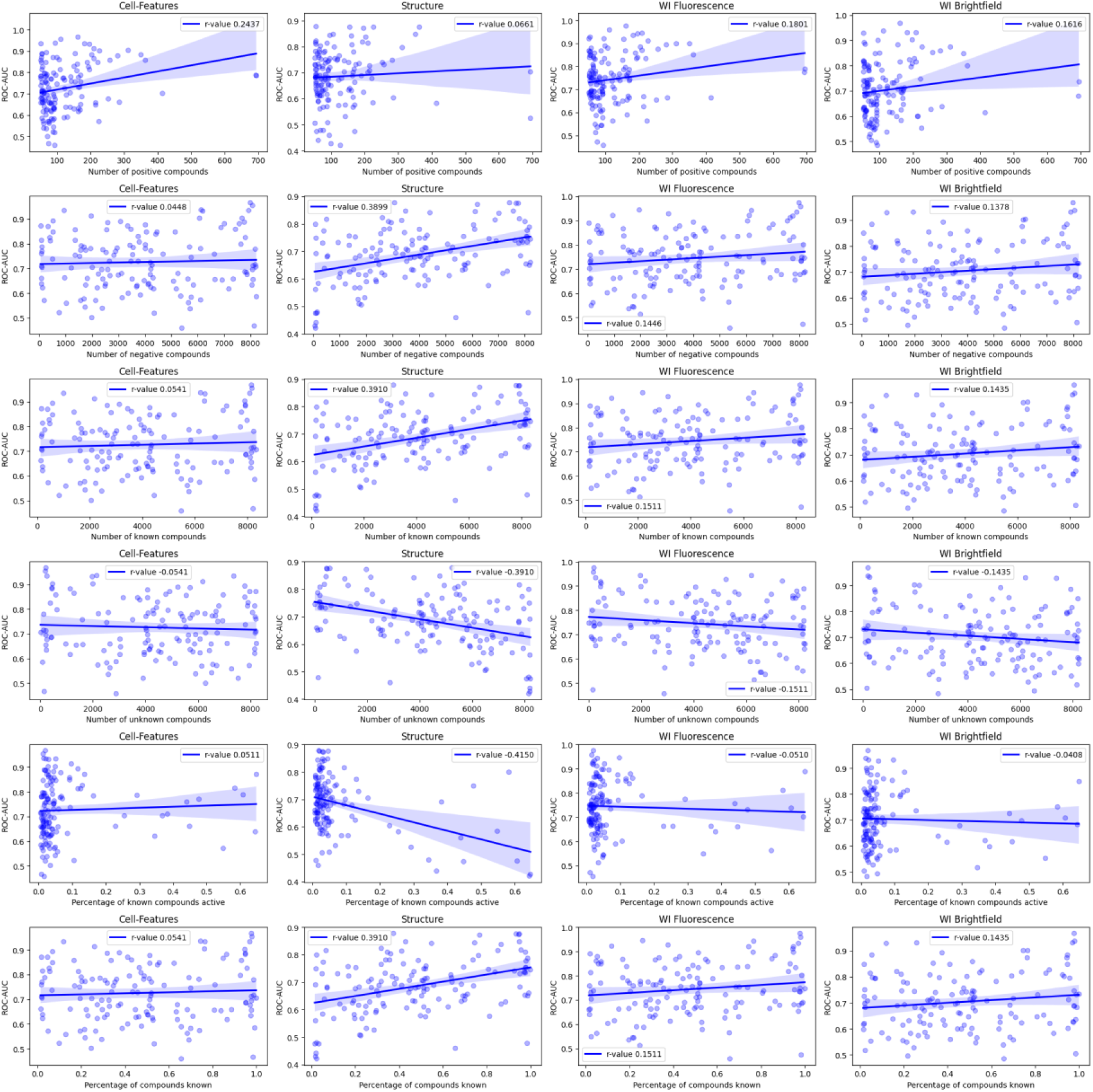
Model performance vs. HTS data availability. Modality-wise performance per assay vs. HTS assay data point availability. Each column represents one input modality, rows represent, 1.) The number of known active compounds, 2.) number of known inactive compounds, 3.) number of compounds with known activity/inactivity, 4.) Number of compounds with unknown activity, 5.) percentage of known compounds being active and 6.) percentage of compounds with known activity readouts.

**Figure 7 – Supplementary figure.**
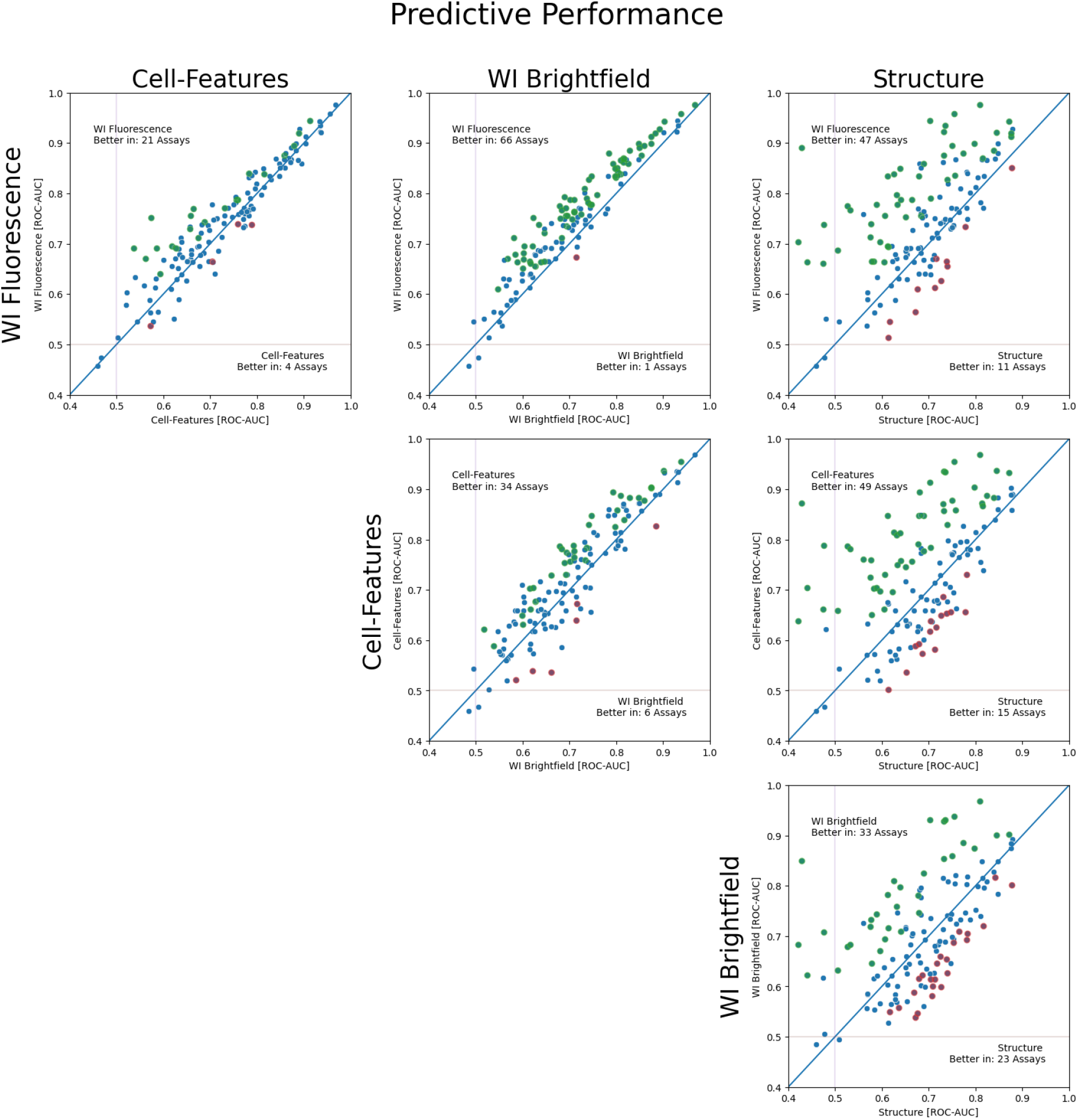
Assay wise performance comparison between Modality types. Comparative performance plot, representing mean assay vise performance for one modality each per axis. Each dot represents an assay, with color representing if there is a significant difference in performance over 6-splits. Green – Modality on y-axis performing better, Red – Modality on x-axis and blue not showing significance. Statistical test using paired Wilcoxon signed-ranked test, with p < 0.05 set as cutoff for use of red/green coloring.

